# Function-oriented microglial transcriptional scores in Alzheimer’s disease: resolving terminal functions beyond activation states

**DOI:** 10.64898/2026.10.14.759278

**Authors:** Alex Tu, Siman Liu

## Abstract

Alzheimer’s disease (AD) staging based on amyloid-β, tau, and neurodegeneration does not capture heterogeneous microglial functions. Existing transcriptomic scores indicate microglial activation or state similarity but do not resolve post-activation functions and are not directly comparable across scales. Herein, we develop a function-oriented framework comprising seven harmful- and seven adaptive-associated gene modules, selected from predefined mechanisms and direct functional evidence. Using SEA-AD microglia/perivascular macrophage nuclei, we compute do-nor–brain-region pseudobulk scores on a fixed P10–P90 reference scale and validate them against matched protein, activity, and tissue endpoints. H3 is related to cytokine/chemokine proteins; H4 and A5 are related to the same isoprostane readout in opposite directions; and A3 is related to dense-core plaque fraction. Benchmark comparison shows modest incremental fit but distinct outcome-specific information. In external GSE146639, eight modules remained FDR-significant, although harmful and adaptive arms are highly collinear; H7 and A2 are replicated in external proteomics. Downsampling, leave-one-gene-out, and alternative scoring analyses supports technical stability, and single-cell reconstruction supported a state-resolved basis. These scores summarize relative engagement of damage-related and adaptive programmes, enabling mechanistic comparison across donors and brain regions, but remain observational, are not intended for clinical decision-making, and require further validation.

## 1. Introduction

Alzheimer’s disease (AD) staging based on amyloid-β pathology, tau pathology, and neurodegeneration does not reflect potential differences in microglial function among similar-appearing patients or across brain regions with comparable macroscopic pathology [1]. Microglia clear plaques and cellular debris [2, 3], metabolize lipids [4], and relieve oxidative stress [5], while also promoting tissue repair [6]. Conversely, they can activate inflammatory pathways [7], phagocytose neural structures [8], injure the neurovascular unit [9], and redistribute toxic cargo [10].

Five major microglial states have been recognized: disease-associated (or activation) states, inflammatory states, lipid-associated states, interferon-related states, and homeostatic states [11–15]. These programs have largely been studied separately, each with distinct gene sets, and their different scoring scales preclude direct comparison [11–15]. State-based scores such as the disease-associated microglia (DAM) signature and the homeostatic signature have been widely applied to AD datasets and have revealed disease-associated microglial responses. However, they primarily indicate whether a response is present rather than what microglia actually do, and they cannot resolve terminal functions such as inflammation, oxidative injury, clearance, or repair. We therefore constructed a function-oriented framework and compared it directly with six microglial benchmark scores in the same SEA-AD donor–region samples. This framework comprises seven harmful-associated and seven adaptive-associated gene groups, selected according to predefined mechanisms and direct functional evidence. The overall harmful and adaptive scores summarize the relative transcriptional engagement of their constituent functional modules. They are primarily intended to reveal differences in microglial functional profiles between donors with similar amyloid and tau burden, or between brain regions with comparable pathological burden. Each individual module can further indicate which terminal mechanism is relatively active, whether inflammation, oxidative injury, vascular injury, clearance, antioxidant defence, or repair. Thus, the framework does not merely determine whether microglia are activated; it determines what microglia predominantly do after activation. An adequately powered study could use this framework to reveal tissue or cognitive damage at similar pathological levels, to assess whether a treatment inhibits pathological but not adaptive functions, and to determine whether human microglial functions in vivo and in vitro are recapitulated in animal, human iPSC-microglial, human organoid, or drug-treated human microglial models. It could also compare fixed-gene, fixed-anchor functional patterns across donors, brain regions, disease stages, cohorts, or experimental systems.

Herein, we validated this framework by examining whether each score agreed with its matched biological readout, whether associations persisted after accounting for local pathology, whether each module reflected its intended mechanism, and whether results were reproducible at the single-cell level. Several modules showed expected biological relationships: H3 was related to cytokine and chemokine proteins; H4 and A5 were related to oxidative-stress measurements in opposite directions; and A3 was related to dense-core plaque fraction. In an external microglial RNA dataset, eight modules remained significant after FDR correction, and H7 and A2 showed expected differences in an external proteomics dataset. Technical robustness analyses further supported stability. These scores therefore provide a function-oriented summary of microglial transcriptional engagement, enabling mechanistic comparisons across donors and brain regions. However, they remain observational, are not intended for clinical decision-making, and require further mechanistic and prospective validation.

## 2. Results

### 2.1 Construction of harmful and adaptive gene groups

To build a function-oriented framework, we first defined 14 gene groups that represent distinct terminal microglial programmes. Based on the previously defined three-level framework, we constructed 14 gene groups linking AD clinical outcomes to tissue-level degeneration and the cellular injuries proximal to these changes. Seven groups were defined as harmful-associated (H1–H7) and seven as adaptive-associated (A1–A7). Each module was biologically paired with one or more related programmes according to its principal function. In the harmful arm, H1 represents immune-tagged neural-material clearance [8]; H2, stressed neural-material removal [16]; H3, inflammatory and cytotoxic mediator release [7]; H4, oxidative-injury generation [17]; H5, neurotoxic kynurenine metabolism [18, 19]; H6, extracellular-matrix and neurovascular injury [9]; and H7, extracellular-vesicle-mediated toxic-cargo spread [10]. In the adaptive arm, A1 represents lipid and myelin-debris handling [4]; A2, intracellular cargo processing and degradation [20]; A3, plaque and debris engagement and processing [3]; A4, antibody-supported clearance signalling [21]; A5, antioxidant defence [5]; A6, homeostatic surveillance and maintenance [22]; and A7, trophic, repair, and remyelination support (Table S1) [6, 23]. These modules formed the basis for all subsequent scoring and validation analyses.

**Table 1.** Harmful-associated and adaptive-associated microglial gene groups and their constituent genes.

| Arm | Gene group | n | Genes |
| --- | --- | --- | --- |
| Harmful | H1 | 4 | CD68, C3, SYK, VAV1 |
| Harmful | H2 | 9 | ADGRG1, CD68, ITGAX, ITGB5, MERTK, P2RY6, SYK, VAV1, TREM2 |
| Harmful | H3 | 10 | CASP1, MAPK14, GSDMD, IL6, IRAK4, MYD88, PTGS2, TRAF6, NLRP3, TNF |
| Harmful | H4 | 3 | LTC4S, RAC1, PLA2G4A |
| Harmful | H5 | 2 | HAAO, KMO |
| Harmful | H6 | 6 | ADAM17, CTSS, VEGFA, CTSD, CTSB, MMP2 |
| Harmful | H7 | 10 | ARRDC1, CD81, HGS, TSG101, SDCBP, SMPD3, STX4, VPS4A, VPS4B, CD9 |
| Adaptive | A1 | 7 | ABCA1, ABCA7, APOC1, LPL, NR1H3, RXRB, SOAT1 |
| Adaptive | A2 | 7 | ATG5, ATP6V1A, BECN1, GUSB, HEXB, LAMP2, CTSL |
| Adaptive | A3 | 8 | AXL, CLEC7A, LPL, LRP1, MARCO, MERTK, MFGE8, ITGAV |
| Adaptive | A4 | 5 | PIK3CD, SYK, LYN, PLCG2, RAC1 |
| Adaptive | A5 | 10 | CAT, CUL3, GPX4, IDH1, SOD1, SQSTM1, TXN, GCLM, TXNRD1, GSR |
| Adaptive | A6 | 3 | MAFB, SALL1, TGFB1 |
| Adaptive | A7 | 10 | IGF1, PDGFB, TIMP1, HGF, FGF2, BMP7, TIMP2, VEGFB, GPNMB, PDGFA |

### 2.2 Harmful and adaptive scores were constructed from microglial expression, although some scoring genes are also expressed in other brain cells

We next asked whether the scoring genes were predominantly expressed in microglia. All harmful and adaptive scores were constructed exclusively from Micro-PVM nuclei; all expression values used for scoring therefore originate from microglia/PVM. Cross-cell-type analyses assessed whether projections in mixed-cell datasets could be attributed specifically to microglia. Lower specificity indicates possible non-microglial contribution. Canonical microglial markers, including CSF1R, P2RY12, CX3CR1, and SPI1, were clearly detected in the scoring population (Figure 1a) [24, 25], supporting its microglial identity. However, functional modules showed different levels of microglial enrichment. H1, H2, H4, and A4 showed the clearest microglial enrichment, as every gene in these modules was expressed at a higher level in microglia than in all evaluated non-microglial classes. H6 showed a similar pattern, although only five of six genes met this condition (**Figure 1b, c**). Genes related to vesicle biology, lipid metabolism, antioxidant defence, and trophic signalling were not exclusively expressed in microglia and were also detected in other cell types. Among the 94 module entries, 49 had higher expression in microglia than in all evaluated non-microglial cell types (Table S2). High broader-expression risk was 40% for H3, 50% for A3, and 90% for A5; H4 had none (**Figure 1d**). Thus, although some genes are shared across cell types, the scores are derived from microglial expression and should be interpreted with the caveat of broader-expression risk.

**Figure 1.**
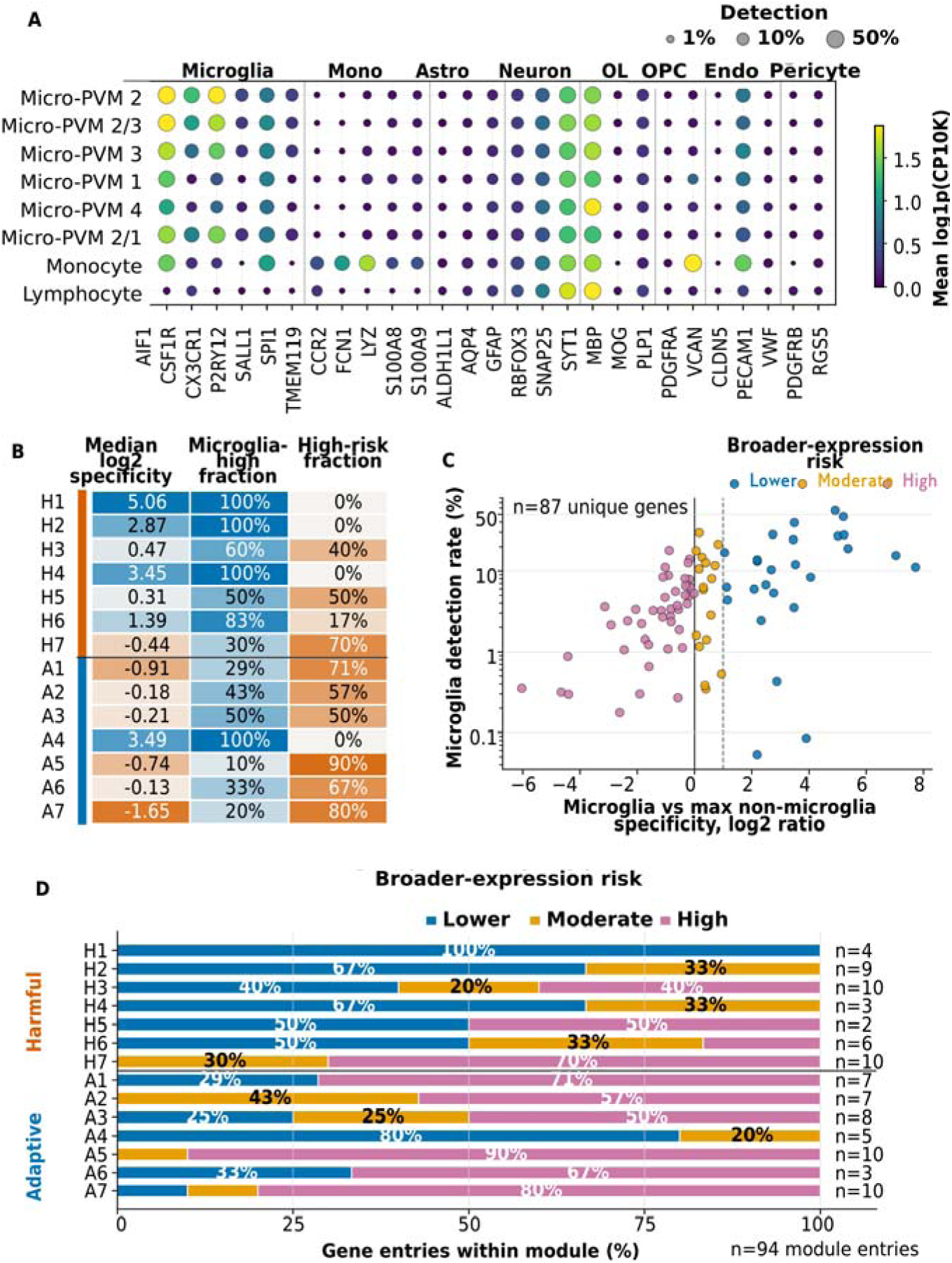
Cross-cell-type expression specificity at the gene and module levels. (a) Marker-expression quality control across SEA-AD Micro-PVM and immune supertypes. Dot size, detection rate; colour, pooled-supertype pseudobulk expression [log1p(CP10K)]. Genes include canonical microglial and non-microglial markers. (b) Cross-cell-type attribution of harmful (H1–H7) and adaptive (A1–A7) modules. Columns show median log2 microglial specificity, percentage of genes with higher microglial expression than all non-microglial classes, and percentage with high broader-expression risk; colour scales differ by column. Exact values are shown in cells. (c) Mean microglial detection rate versus cross-cell-type specificity for 87 unique scoring genes. Specificity is the log2 ratio of microglial expression to the maximum non-microglial expression; detection rate is on a log scale. (d) Broader-expression risk across 94 gene–module entries. Lower risk, ≥2-fold higher microglial expression; moderate, higher but <2-fold; high, not exceeding non-microglial expression. n, entries per module; genes in multiple modules were counted in each. Alt text: Microglial expression specificity varies across 14 modules and non-microglial cell types.

### 2.3 RNA signals of selected modules were related to their corresponding product/activity readouts and tissue endpoints

To assess whether module scores reflect their intended functions, we examined their relationships with matched product/activity and tissue endpoints. In the same donor and brain region, 12 independent module–readout associations were available from eight modules: five RNA–product/activity relationships, five RNA–tissue-endpoint relationships, and two shared oxidative-stress readouts (H4 and A5 isoprostane, counted once; Figure 2A and Table S3). H4 and A5 use the same isoprostane analyte across the product/activity and tissue-endpoint layers; to avoid pseudoreplication, they were removed from the protein/activity and tissue-endpoint blocks and are displayed only once in the shared oxidative-stress readout block. Coefficients were aligned with the expected module function, so a positive value follows the expected harmful or adaptive direction. For product/activity readouts, H3 showed the strongest relationship with the cytokine/chemokine protein index (β = 0.181, 95% CI 0.068–0.293, P = 0.0016, q = 0.0097). H6 and A6 reached nominal significance. A3 and A7 followed expected directions but were not statistically significant. For tissue endpoints, A3 was related to a higher dense-core plaque fraction (β = 0.221, 95% CI 0.035–0.406, P = 0.0197, q = 0.0197), consistent with microglial plaque engagement and organization. H3, H7, and A6 generally aligned with expected tissue-endpoint directions, although none were statistically significant, and A7 did not show the expected trend. In the shared oxidative-stress readout block, H4 (β = 0.244, P = 0.0132, q_shared = 0.0467) and A5 (β = 0.254, P = 0.0156, q_shared = 0.0467) were both related to the same isoprostane analyte in the expected opposite directions. This shared readout was counted once and does not constitute two independent evidence chains or an independent RNA–product–endpoint chain. Residual plots for four independent relationships are shown in Figure 2B–E. These results support several module–readout relationships, with H3, H4, A5, and A3 showing the clearest expected associations.

**Figure 2.**
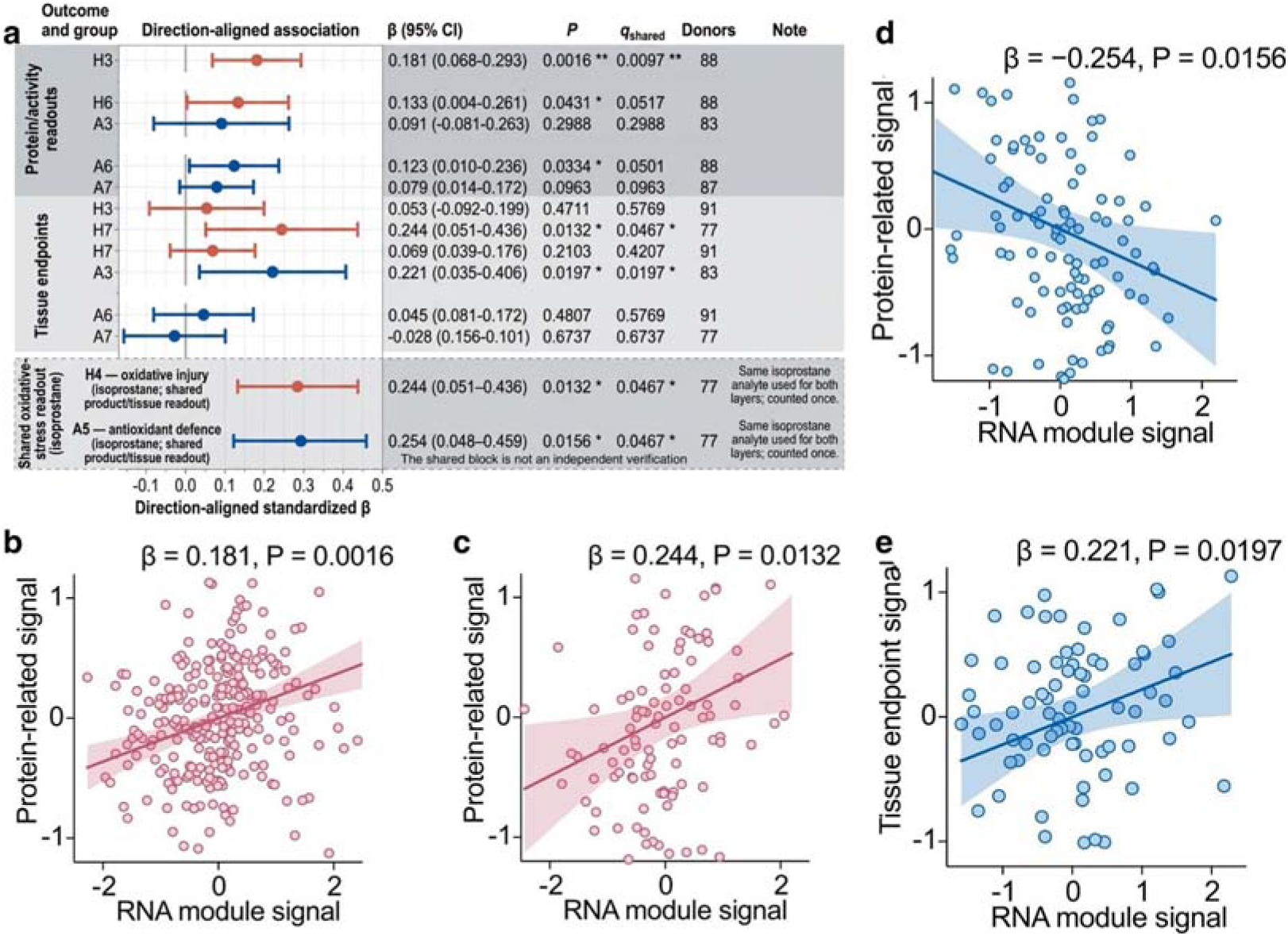
Relationships between module RNA scores and matched product/activity, tissue-endpoint, and shared oxidative-stress readouts. (a) Direction-aligned associations for estimable module–readout pairs. (b–e) Adjusted donor–region residual scatter plots for independent readouts: (b) H3–cytokine/chemokine protein index; (c) H4–isoprostane shared oxidative-stress readout; (d) A5–isoprostane shared oxidative-stress readout; (e) A3–dense-core plaque fraction. Symbols: *P or q < 0.05; **P or q < 0.01; ***P or q < 0.001.

### 2.4 Diagonal specificity of matched module–endpoint relationships

We next asked whether these relationships were specific to the matched module rather than reflecting general disease severity. Diagonal specificity determines whether a target module has a stronger relationship with its corresponding endpoint than with the other 13 modules. Only H3, H7, A3, A6, and A7 had eligible endpoints; thus, 11 relationships were tested (Tables S3, S4). Among these, nine showed the expected direction (Figure 3), and eight ranked within the top four among all 14 modules. The strongest relationship was between A3 and dense-core plaque fraction; A3 ranked first for this endpoint and remained significant after FDR correction. A6 ranked third or fourth for each of the three IBA1 endpoints. A7 ranked third for both NeuN endpoints, and both relationships reached nominal P < 0.05. These results mainly support the matched relationships of A3, A6, and A7 with their corresponding endpoints. H3 and H7 showed weak relationships with directions opposite to expectation (Table S4). A3, A6, and A7 showed the strongest diagonal specificity, whereas H3 and H7 were non-specific.

**Figure 3.**
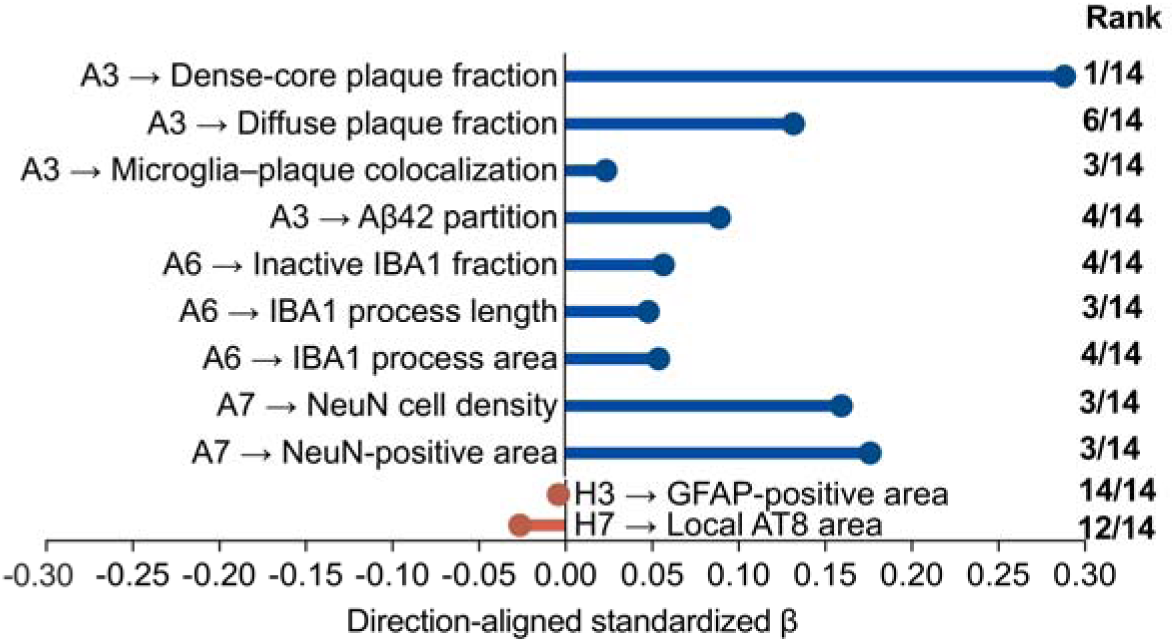
Relative specificity of matched module–endpoint relationships. Direction-aligned standardized β from two-sided donor-clustered linear models of donor–region observations. Ranks indicate each target module’s position among all 14 modules based on absolute standardized β. BH correction was applied across 154 models. Harmful modules are shown in red and adaptive modules in blue. Exact P values, FDR q values, donor counts, and ranks are provided in Table S4. Alt text: Direction and rank of 11 matched module–endpoint relationships; A3, A6, and A7 rank highly for several endpoints.

### 2.5 Comparison with microglial benchmark scores

We then compared the modules with established microglial benchmark scores to determine whether they provide independent information. For tau and amyloid analyses, we included 497 donor–region observations from 84 donors. For NeuN, 156 observations from the same donors were included. We first examined the overlap between the 14 function-oriented modules and six microglial benchmark scores (Figure 4a). Among all 84 module–benchmark pairs, the median absolute Spearman correlation was 0.180, and 61 pairs showed a correlation lower than 0.30, indicating that the modules were not simple reproductions of existing benchmark scores. After each benchmark was considered separately, 19 module terms among the 252 terms remained FDR-significant. In contrast, when the related module was included, only 7 benchmark terms among the same 252 terms remained FDR-significant. The median partial R² added by the modules was 0.0096, compared with 0.0047 for the benchmark scores, and the full model had a lower AIC than the benchmark-only model in 157 of 252 comparisons (Table S5). We next performed donor-grouped cross-validation to assess incremental model fit for tau, amyloid, and NeuN (Figure 4b). Compared with benchmark-only models, full models reduced RMSE in 112 of 252 comparisons. However, compared with module-only models, RMSE was reduced in only 54 of 252 comparisons. These results indicate that full models provided modest incremental fit over benchmark-only models but did not consistently outperform module-only models. Therefore, the value of the framework lies primarily in mechanistic decomposition rather than improved prediction. To determine whether the modules captured outcome-specific variance beyond benchmark scores, we examined full-model coefficients for the clearest relationships (Figure 4c). A3 retained a significant association with tau (module P = 0.012), A1 with amyloid (module P = 0.012), and H6 with NeuN (module P = 0.02), whereas the corresponding benchmark terms were not significant (P = 0.34, 0.34, and 0.37, respectively). These results indicate that selected function-oriented modules capture outcome-related information not fully represented by individual microglial benchmark scores. Together, these results indicate that the framework provides modest incremental fit but distinct outcome-specific information, supporting mechanistic decomposition over prediction.

**Figure 4.**
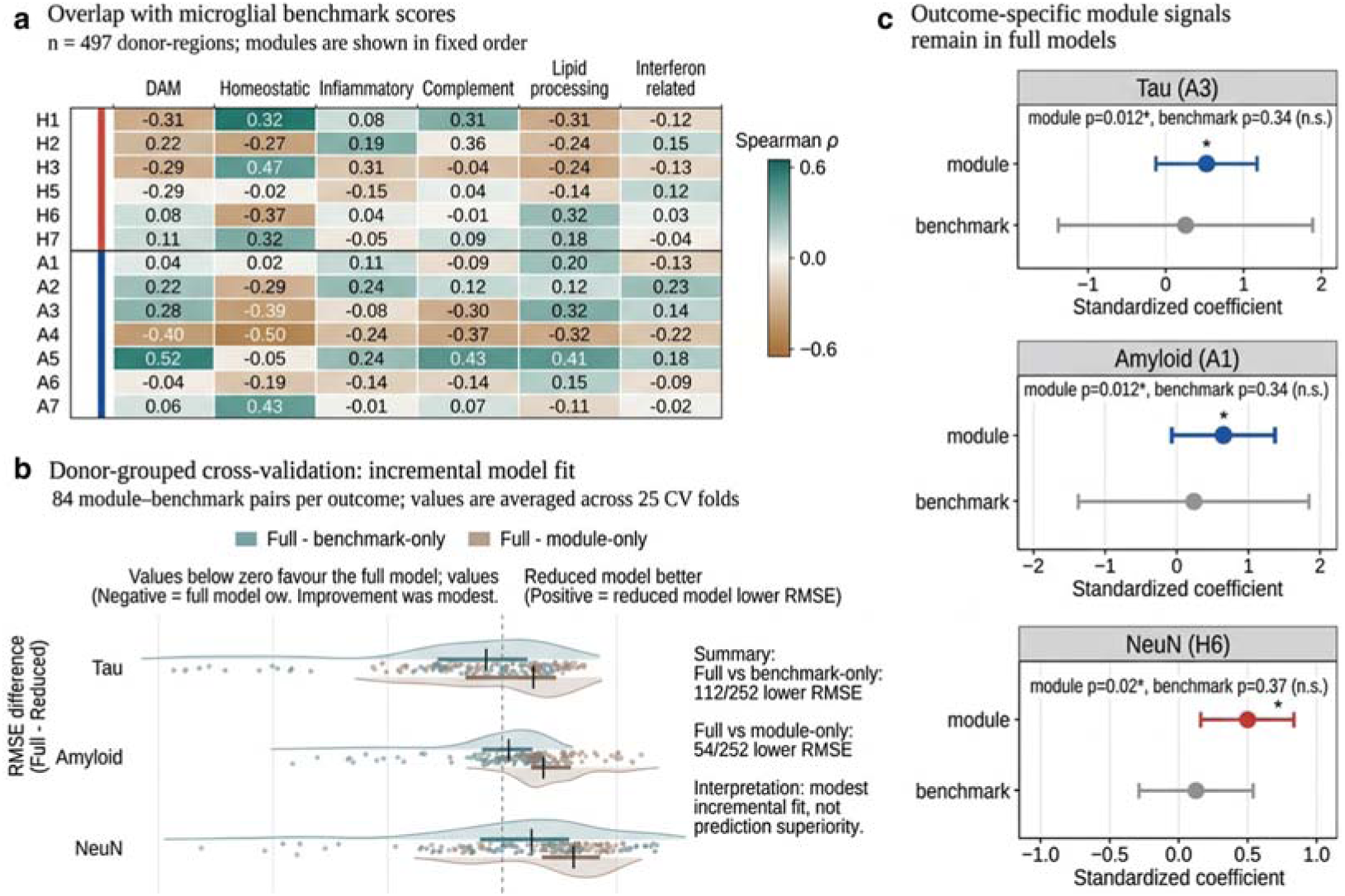
Comparison with microglial benchmark scores and outcome-specific signals in full models. (a) Spearman correlations between the 14 modules and six microglial benchmark scores in 497 SEA-AD donor–region observations. Red and blue side bars indicate harmful- and adaptive-associated modules, respectively.(b) Donor-grouped cross-validation (5 repeats of 5-fold CV, 84 donors) for tau, amyloid, and NeuN. Points represent RMSE differences: full model minus benchmark-only (teal) or full model minus module-only (taupe). Negative values indicate lower RMSE for the full model. Horizontal bars show interquartile ranges; black ticks show medians. Full models reduced RMSE in 112/252 and 54/252 comparisons, respectively.(c) Full-model standardized coefficients for selected modules. A3 (tau), A1 (amyloid), and H6 (NeuN) retain significant associations (module P values shown; benchmark P values are n.s.). Points represent coefficients; error bars represent 95% CIs with donor-clustered robust standard errors. Harmful modules are shown in red, adaptive modules in blue, and benchmark scores in grey. n.s. = not significant.

### 2.6 Module signals could be reproduced in internal splits and external datasets

To test reproducibility, we examined internal split stability and external RNA and protein datasets. The module–readout relationships of the internal discovery and locked-holdout sets were compared to assess stability (Figure 5a). Expected directionality was maintained in the majority of relationships across both sets. In the Allen-based analysis, nine of ten relationships were directionally consistent across all splits. Similarly, both SEA-AD A3 readouts preserved expected directions across all splits. Overall, 66.4% of Allen target–split combinations followed the expected direction. In A3 analyses, the expected direction was preserved in 64% of Aβ42 partitions and 72% of dense-core-plaque analyses. Two relationships showed reversed directions: A6 cytokine activity in the discovery set and A7 BDNF in the locked-holdout set. For external RNA replication, Smart-seq2 data from GSE146639 were used, comprising FACS-isolated human CD11B+/CD45+ microglia aggregated at the donor level. All seven harmful-associated modules exhibited positive coefficients, whereas all seven adaptive-associated modules exhibited negative coefficients. Eight modules remained significant after FDR correction: H1, H4, H5, H7, A1, A2, A3, and A5 (Figure 5b). The strongest relationships included H4 (β = 0.81, q = 0.0014), H7 (β = 0.52, q = 0.0012), A1 (β = −1.12, q = 0.0068), and A5 (β = −0.29, q < 0.001). This pattern persisted across all 14 modules after covariate adjustments and sensitivity analyses; eleven modules remained stable in leave-one-donor-out analyses, while H3, H6, and A7 were donor-sensitive. To explicitly test inter-arm independence in the external dataset, we correlated donor-level Harmful and Adaptive total scores (Figure 5c). The two arms were highly collinear (Spearman ρ = −0.96, P < 0.001, n = 14 donors). This suggests that the functional dichotomy may be compressed in acute post-mortem FACS-isolated microglia compared with the SEA-AD nuclear dataset, underscoring the need for cautious interpretation of arm independence across different experimental platforms. At the protein level, H7 levels were higher in AD and A2 levels were lower in AD compared with controls (H7: β = 0.73, q = 0.026; A2: β = −0.40, q = 0.026; Figure 5d). Relationships with asymptomatic AD also followed expected directions but did not reach statistical significance after FDR correction. Collectively, these results demonstrate internal stability and external directional reproducibility of the module signals. These analyses demonstrate internal stability and external directional reproducibility, while highlighting the need for cautious interpretation of arm independence in external datasets.

**Figure 5.**
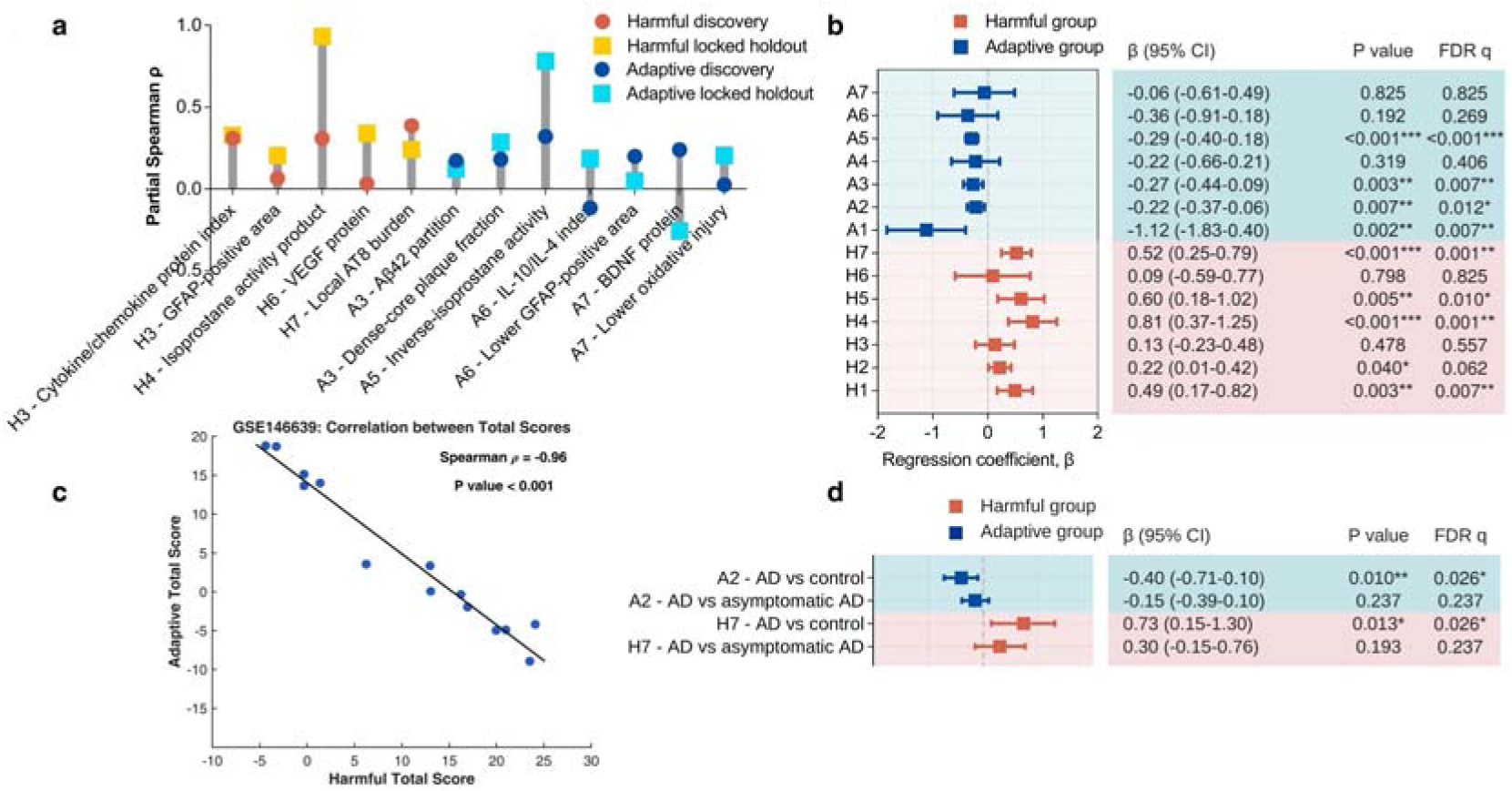
Reproducibility of harmful and adaptive module signals in internal and external datasets. (a) Direction-aligned partial Spearman correlations between discovery and locked-holdout sets across ten Allen readouts and two SEA-AD A3 readouts. Lines connect estimates from the same readout. A positive value indicates the expected direction. Descriptive split-stability comparison only. (b) External RNA validation in donor-aggregated GSE146639 microglia (n = 14 donors). Points represent age- and sex-adjusted regression coefficients with HC3 robust standard errors; bars represent 95% CIs. All harmful modules (red) show positive coefficients, while all adaptive modules (blue) show negative coefficients. Benjamini-Hochberg correction was applied across 14 module tests. (c) Donor-level correlation between Harmful and Adaptive total scores in GSE146639 (n = 14 donors). Each point represents one donor. The two arms were highly negatively correlated (Spearman ρ = -0.96, P < 0.001), indicating that inter-arm independence was not preserved in this external dataset. This suggests that the functional dichotomy may be compressed in FACS-isolated microglia compared to the SEA-AD nuclear dataset. (d) External protein validation (E-PROT-51) for H7 and A2 in dorsolateral prefrontal cortex. Models are age- and sex-adjusted with HC3 robust standard errors (n = 51 for AD vs. control; n = 54 for AD vs. asymptomatic AD). Benjamini-Hochberg correction was applied across four comparisons. *P or q < 0.05; **P or q < 0.01; ***P or q < 0.001. Alt text: Internal and external analyses compare module signals across data splits, an independent microglial RNA dataset (including arm-level collinearity), and an external protein dataset.

### 2.7 Single-cell-state reconstruction was reproducible across non-overlapping nuclei

We next asked whether the scores could be reconstructed from single-cell states and whether this reconstruction was reproducible. Direct reconstruction showed expected agreement between observed and reconstructed scores (Figure S4), prompting us to test cross-half reproducibility across non-overlapping nuclei. Across 100 random within-donor-region splits, the observed score in one half remained correlated with the complementary-half reconstruction from the other half. The median cross-half rho was 0.671 for the harmful score and 0.626 for the adaptive score, reaching 98.1% and 97.4% of the corresponding split-half reliability ceilings, respectively. Thus, the state-based score structure was reproducible across non-overlapping nuclei within the same RNA dataset (Table S7). We then examined whether module scores were driven by within-state expression intensity or state abundance (Figure 6a, b). Module scores showed stronger positive correlations with within-state intensity, particularly in the dominant PVM2 state (median rho = 0.874), whereas correlations with state fractions remained close to zero overall (median rho = −0.007). These same-RNA compositional analyses are descriptive and do not estimate causal state contributions. These results support a single-cell-state basis for the scores but do not establish causal state contributions.

**Figure 6.**
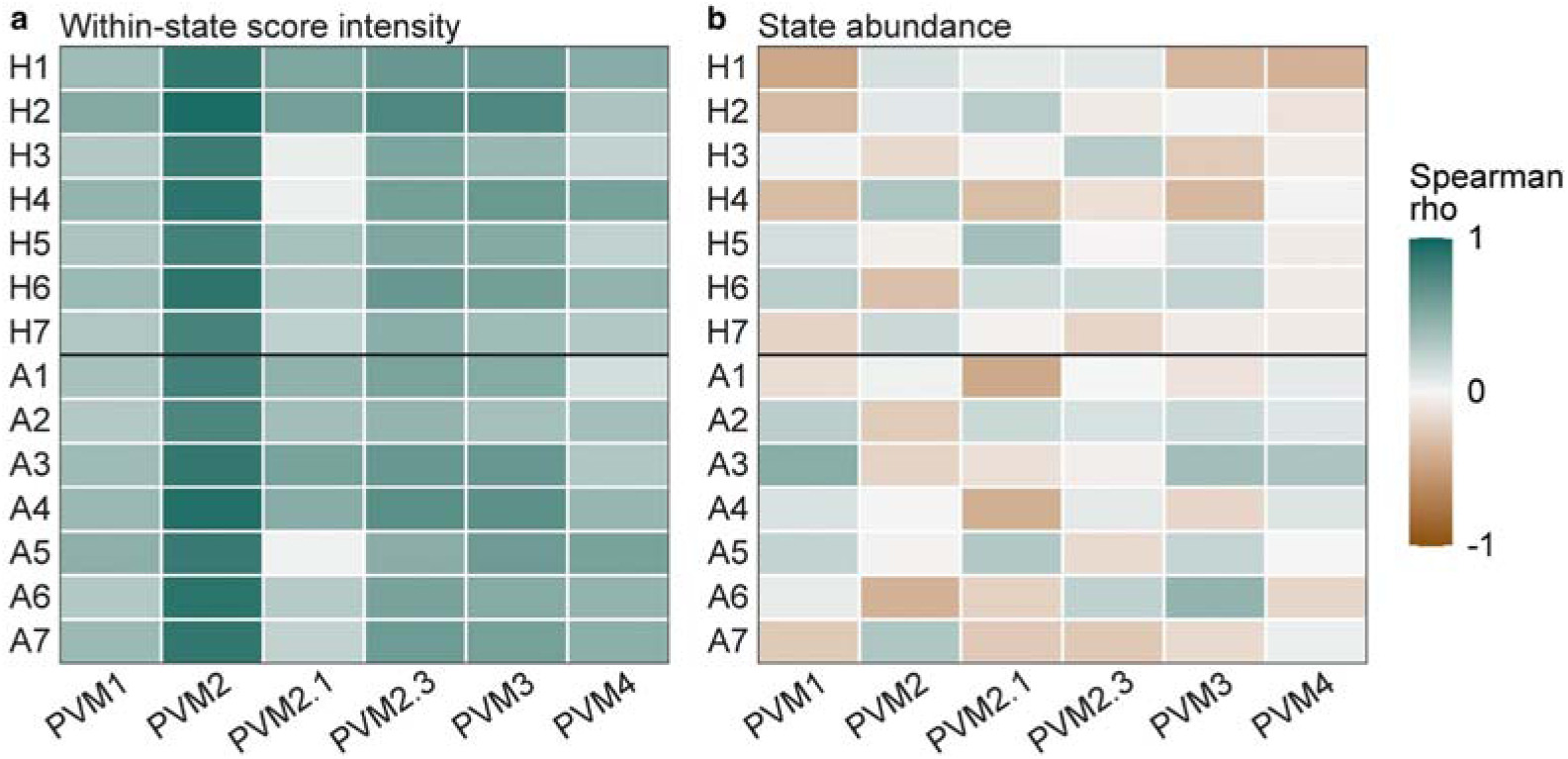
Module scores correspond more closely to within-state expression intensity than to state abundance. (a) Within-state expression intensity heatmap. (b) State fraction heatmap. State-specific analyses included donor regions with at least 10 nuclei in the indicated state (PVM1 n=294; PVM2 n=528; PVM2.1 n=31; PVM2.3 n=505; PVM3 n=491; PVM4 n=31).

### 2.8 Technical robustness of harmful and adaptive scores

To determine whether harmful and adaptive scores remained stable under different technical conditions, we examined the effects of cell number, individual genes, and scoring method. For every eligible donor–region, we randomly retained 200 microglial nuclei and repeated this process 100 times. Harmful and adaptive scores calculated from downsampled data remained close to scores calculated from full data. Spearman correlations between full-data and downsampled scores were 0.981 for harmful and 0.976 for adaptive. Single-run ICC(1,1) values were 0.928 and 0.931 (Figure 7a), respectively. For all 14 modules, correlations after downsampling were higher than 0.961 (Figure 7b), with ICC values ranging from 0.833 to 0.985. H3, H7, and A2 had ICC values lower than 0.90; however, donor rankings of these modules still correlated highly with rankings from full-data scores. To test whether the overall harmful or adaptive arm was driven by a single gene, we removed every gene separately and recalculated scores. Among the 94 gene-entry deletions, the lowest correlation at the arm level was still 0.915. Hence, the overall arms are not driven by one single gene. At the module level, most gene deletions caused only small changes (median correlation 0.964). The clearest local changes appeared in several small modules: H5 after KMO removal, A6 after MAFB removal, and A2 after HEXB removal (Figure 7c).

**Figure 7.**
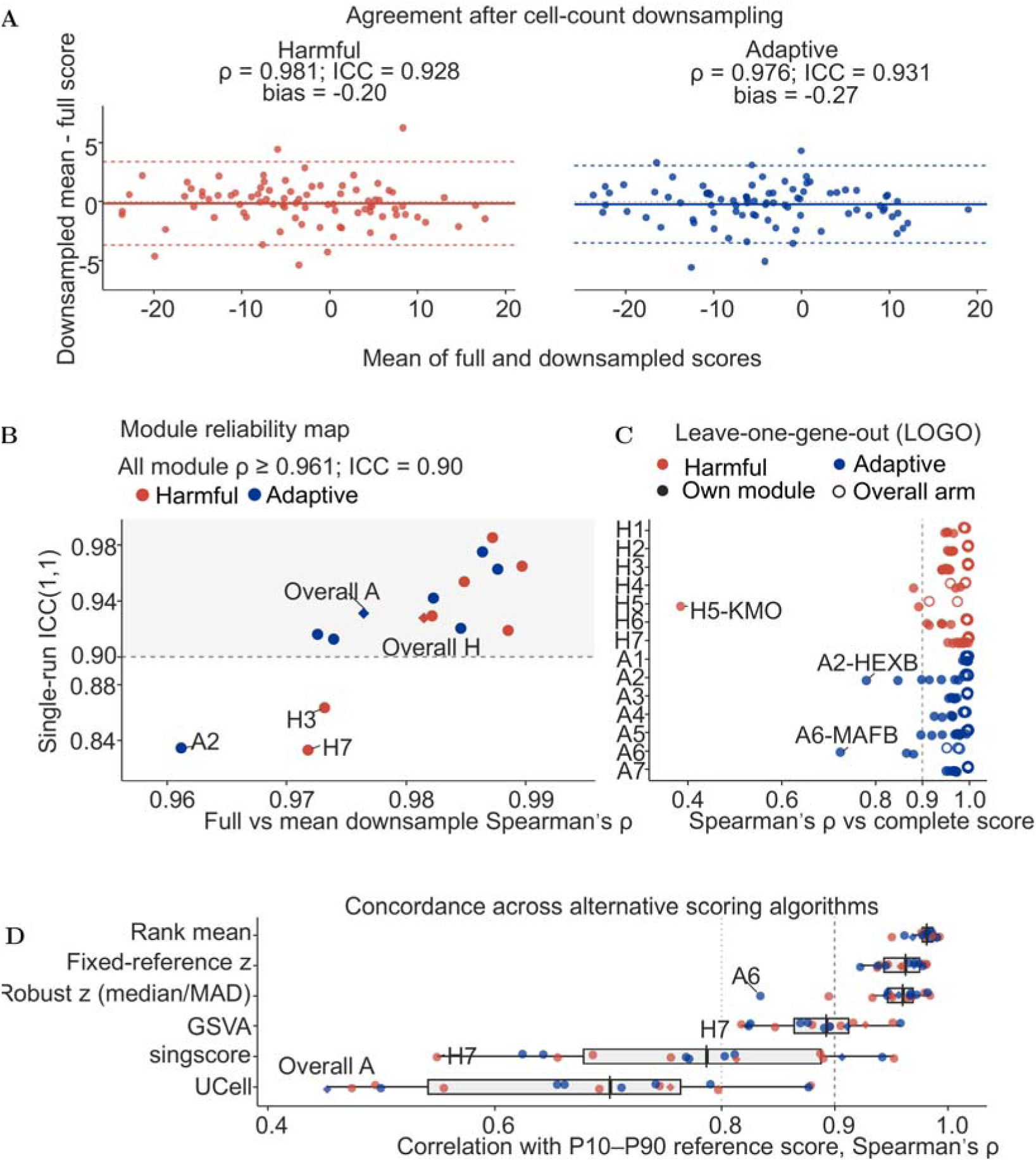
Technical robustness of the harmful and adaptive scores. (a) Bland–Altman plots comparing full-data donor scores with mean scores from 100 cell-count downsampling runs (200 Micro-PVM nuclei per donor–region, reaggregated at donor level; n = 84 donors). Grey dotted line, zero difference; solid line, mean bias; dashed lines, 95% limits of agreement. ρ, Spearman correlation between full and downsampled scores; ICC, single-run ICC(1,1). (b) Module reliability: x-axis, Spearman’s ρ between full-data and downsampled scores; y-axis, single-run ICC(1,1). Dashed line, ICC = 0.90. Circles, modules; diamonds, overall arms. (c) Exact leave-one-gene-out analysis across 94 gene entries. Points show correlation between the complete score and the score after removing one gene entry. Filled circles, module directly losing the gene; open circles, corresponding overall arm. Dashed line, ρ = 0.90. (d) Six alternative scoring algorithms compared with the P10–P90 reference score. Boxes, middle 50% of correlations; whiskers, 1.5× interquartile range; black ticks, medians. Vertical lines, ρ = 0.80 and 0.90. Harmful scores in red; adaptive scores in blue. These are descriptive technical-sensitivity analyses, not tests of biological validity or causality. Alt text: Downsampling, gene removal, and alternative scoring methods test technical stability of module and overall scores.

The exact scoring method could also influence donor rankings. We used several alternative scoring methods and compared their results with the P10–P90 reference score. Rank-based scores and fixed-reference continuous scores produced similar donor rankings. Median correlations were 0.982 for rank mean, 0.963 for fixed-reference z-score, and 0.960 for robust z-score (Figure 7d). GSVA showed moderate agreement with the reference score, while singscore and UCell showed weaker agreement. According to these results, the main harmful and adaptive signals are technically stable. However, several small modules can be more sensitive to technical changes. Specifically, modules with fewer than four genes (H4, H5, and A6) showed higher deletion sensitivity in leave-one-gene-out analyses. Therefore, interpretation of these specific sub-scores should be module-level only, and their constituent genes should not be used as independent biomarkers without further validation (Table S8). Collectively, these analyses establish a technically stable framework, with the caveat that small modules should be interpreted at the module level only.

## 3. Discussion

Our validation was designed to determine what evidence should be observed if a harmful or adaptive score reflects a microglial function. Such evidence should originate from microglia, relate to the expected functional output, persist after accounting for alternative explanations, show the expected relationship with the corresponding endpoint, and be reproducible and stable. The scores were constructed using SEA-AD Micro-PVM nuclei. Genes were selected because their encoded proteins directly participate in predefined terminal mechanisms, not because their expression correlated with AD severity. We also examined expression of these genes in other brain cell types to assess whether non-microglial expression could affect interpretation. This analysis showed that not every scoring gene is expressed exclusively in microglia, which is most relevant for H3, A3, and A5, with high broader-expression risk of 40%, 50%, and 90%, respectively; H4 had none.

We examined whether a higher RNA score was accompanied by the product or tissue change expected from this function in matched donor–region samples. H3 was related to cytokine and chemokine proteins. H4 and A5 were both related to the same isoprostane readout in expected opposite directions; this shared oxidative-stress measurement was counted once and does not constitute two independent validations. A3 was related to dense-core plaque fraction, consistent with microglial plaque engagement and organization [3, 26]. Residual plots show that observed relationships are not driven by a single outlier. Importantly, however, these bulk-RNA data show tissue-level correspondence with similar readouts, not microglia-specific functional validation. To ensure that an observed module-specific relationship was not simply amyloid or tau burden, we compared relationships for each module to each of the other 13. Modules expected to associate with increased dense-core plaque fraction ranked near the top (especially A3). That many modules were specific for their mechanism beyond disease severity suggests associations with H3 and H7 may show non-specificity.

Benchmarking revealed small overlaps and partial, outcome-specific non-redundancy between A3–tau, A1–amyloid, and H6–NeuN. These results suggest that functional decomposition of different biomarkers is preferable to combined models when combining complements does not improve prediction. Comparisons between discovery and locked-holdout sets reflected persistent external microglial RNA and protein signals, although donor-specific influence remained due to the reduced number of donors. Evidence from external protein data further affirmed these findings. Our state inference from single cells showed stronger correlations between scores and transcriptional activity than with abundance of microglial states. We controlled for cell number in downsampling analysis and for the effect of a particular gene with leave-one-gene-out analysis. Other scoring systems were attempted with similar results, which cannot be attributed to cell number, a single gene, or a single scoring algorithm.

Together, these analyses provide observational support strongest at the module level, but overall these scores are summary statistics, not validated functional measures. Furthermore, being only an observational study, one cannot infer that changing one of these scores will change how microglia and tissue respond to injury. Not all modules had matching endpoints: some were weakly related, and others inversely related. The number of external samples was small. H4 and A5 used the same isoprostane measurement; this shared readout was treated as a single oxidative-stress measurement and does not provide independent verification of two separate biological layers. Protein-level replication involved only H7 and A2. Direct reconstruction used identical RNA data, and complementary-half reconstruction used non-overlapping nuclei in the RNA data; neither analysis establishes functional validity. The tests were performed in SEA-AD isolation; post-mortem scores are not currently linked with prognosis or treatment outcome. Finally, moderate-to-low correlations with GSVA and singscore indicate that our scores are not algorithm-agnostic. The fixed-reference scaling was deliberately chosen to anchor expression to a biologically meaningful population distribution; however, users should be aware that applying alternative single-sample scoring methods without this anchor may dilute module signals.

## 4. Materials and Methods

### 4.1 Selection of harmful-associated and adaptive-associated microglial genes

Genes were selected using a mechanism-based eligibility procedure. Harmful-associated and adaptive-associated modules were constructed independently. For each module, we first defined the microglial function to be represented and divided this function into the biological steps required to complete it. Candidate genes were then identified through PubMed searches. UniProt [27], Gene Ontology [28], and Reactome [29] were used to search for additional terms and candidates. Reviews and database annotations were only used to generate these candidates and were not used to directly determine whether a candidate could enter the final gene set. A candidate was retained only when its function was supported by primary experimental evidence.

A gene could enter the final gene set only when the protein encoded by this gene directly completed one necessary step of the mechanism. If the protein did not directly complete this step, it had to be a necessary proximal controller that directly controlled this step. Thus, its manipulation should lead to a change in either the tissue-specific outcome or the product in the organism. The manipulation was accepted to have a deleterious or adaptive effect only when consistently reported in the biological context described in the article and when the direction of the effect was consistent with the previously defined role. We preferred evidence from human or rodent microglia, iPSC-derived microglia, or other CNS myeloid cells. We also considered other cell types if they performed the relevant biochemical activity directly and there was independent evidence that the activity was performed by microglia. We did not consider genes that were only differentially expressed or co-expressed, because these activities alone were insufficient to define microglia. Genes that only appeared in pathway enrichment or disease association analyses were also not sufficient. Genes used only as markers of a cell state were still not sufficient.

After carrying out all the necessary steps, redundant genes were removed unless they resulted from different necessary steps or different mechanism pathways, except where they were required members of one functional complex. If a gene was present in more than one module, the evidence for each gene–module relationship was assessed separately. When data were conflicting or dependent on the biological context, this was indicated and considered in the decision. Human gene symbols were standardized to HGNC nomenclature [30]. Literature-selected genes were then checked in SEA-AD Micro-PVM data [31]. Genes that did not appear in the expression matrix, or were detected in fewer than 1% of Micro-PVM nuclei, were removed. This final filtering step was not used to determine whether a gene was harmful- or adaptive-associated; it was only used to ensure that the gene could be measured reliably.

### 4.2 Calculation of harmful and adaptive scores

After separating samples by donor and brain region, we calculated scores at the donor–brain-region level. Raw counts from all Micro-PVM nuclei of the same donor and brain region were summed to generate one pseudobulk sample. This avoided treating nuclei from the same specimen as independent samples and causing pseudoreplication [32, 33]. Counts were normalized to CP10K (counts per 10,000 UMIs), and normalized expression was transformed using log(1 + CP10K).

Because genes have different expression ranges, each gene was placed on a fixed SEA-AD reference scale. The gene-specific 10th and 90th percentiles represented 0 and 100, respectively. Values between these anchors were transformed by linear interpolation, and values outside the range were limited to 0 or 100. The anchors were fixed and were not recalculated in later SEA-AD analyses.

For each gene group, scaled gene values were combined using fixed gene weights. Gene weights were derived from direction-aligned gene–endpoint associations in development evidence sources and converted from within-module percentile ranks (Table S1). If some genes were unavailable, the remaining weights were renormalized over the observed genes. A module was set to missing only when the dataset-specific coverage threshold was not met. To reduce signals caused by shared biology, each module score incorporated a fixed biologically paired programme:

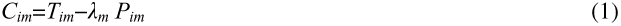

Here, *T_im_* represents the fixed-weight target-programme component, *P_im_* represents the mean fixed-weight component of the biologically paired programme(s), and λ*_m_* controls the paired contribution. The resulting module score represents the integrated transcriptional signal of the target programme in relation to its prespecified biological pairing. The biologically paired programmes were not assumed to be strict biological opposites (Table S1). We calculated the mean of the seven harmful module scores to produce one overall harmful score, and the mean of the seven adaptive module scores to produce one overall adaptive score. Negative values were retained because no clipping was applied. Complete score definitions and parameters are provided in Table S1; the same scoring configuration was used in all validation and robustness analyses. External RNA results are in Table S6.

### 4.3 Validation and robustness analyses

Unless otherwise stated, analyses used donor–brain-region observations. Since several brain regions could originate from one donor, these observations should not be considered completely independent. To avoid treating these regions as independent observations, we used donor-clustered robust standard errors [34]. Continuous variables were standardized before testing. Unless otherwise stated, tests were two-sided. For Figure 2, Benjamini–Hochberg correction used separate Allen product/activity (6 tests), Allen tissue-endpoint (6 tests), SEA-AD A3 product/activity (1 test), and SEA-AD A3 tissue-endpoint (1 test) families (Table S3) [35]. Biological correspondence was evaluated mainly at the gene-group level because the available protein, activity, and tissue readouts were linked to specific mechanisms. The overall harmful and adaptive scores were evaluated as integrated summaries of their constituent gene-group programmes.

#### 4.3.1 Cross-cell-type expression attribution

To determine whether the scoring genes were mostly expressed in microglia, we summarized SEA-AD detection and expression rates for each Micro-PVM and other classes. We then compared the average Micro-PVM expression of each scoring gene with that gene’s maximum average expression across the other classes. The difference between these two values was used to measure the microglial expression specificity of the scoring gene. Specificity was calculated for all genes, and module-level results were summarized by taking the median for each of the 14 gene groups. We also calculated the proportion of genes whose expression peaked in Micro-PVM and the risk that other tissues had higher expression levels (Table S2).

#### 4.3.2 Matching selected readouts and comparison models

We focused on protein/assay readout information available in cohort databases to determine potential readouts and tissue endpoints for each gene group. Most of this information came from the Allen Aging, Dementia and TBI resource and SEA-AD [31, 36]. Readouts had to be available in these resources, biologically related to the gene set being studied, and measured in the required number of samples. This resulted in 377 observations from 107 Allen donors across four brain regions, with bulk RNA expression values log2(normalized FPKM + 1). Expression was z-score normalized by region and projected onto predefined gene groups; associations with bulk RNA were interpreted as bulk-tissue correspondence rather than microglia-specific functional validation. Cell-type specificity of these associations is shown in Table S2. From this, 12 independent module–readout associations were inferred from eight gene groups: five RNA-to-product/activity relationships, five RNA-to-tissue-endpoint relationships, and two shared oxidative-stress readouts (H4 and A5 isoprostane, counted once). For each relationship, the readout signal was regressed against the corresponding gene-group score.

Models further included all available local amyloid and tau measures, as well as demographic, technical, and microglial covariates. Coefficients were aligned such that a positive coefficient indicated that the result was in the hypothesized harmful or adaptive direction (expected direction, defined a priori). The chosen distributions were assessed by six adjusted residual plots. These plots were not treated as new statistical tests (Table S3 and Figure S1–S3).

#### 4.3.3 Diagonal specificity

Rather than assuming that an endpoint was specific to its target group only because one significant relationship was found, we performed the diagonal-specificity analysis. This analysis compared the relationship of the matched endpoint with its target gene group and the other 13 groups. An endpoint could be included only when it was measured in the same donor and the same brain region and was matched with its target group before statistical testing. All groups were analysed using the same model structure. The endpoint could not be calculated from genes included in its target score because this would cause a direct overlap between the score and the endpoint. Eleven relationships from H3, H7, A3, A6, and A7 met these criteria. For every endpoint, we tested all 14 gene groups with the same covariates. We ranked the absolute standardized coefficient of the target group among the coefficients from all 14 groups (Table S4).

#### 4.3.4 Comparison with microglial benchmark scores

We compared H1–H7 and A1–A7 with six microglial benchmark scores. DAM and homeostatic scores were literature-informed [11, 15]; the other four were curated functional benchmarks. Gene lists and scoring rules are provided in Table S5. For tau, amyloid, and NeuN, we constructed the module-only model, the benchmark-only model, and the full model. The same observations and covariates were used in all models. Partial R² and AIC were used as descriptive measures of incremental model fit [37]. For terms in the full model, we used robust standard errors clustered by donor. Benjamini–Hochberg correction was performed separately within each outcome family. We also conducted repeated five-fold cross-validation according to donor groups, and RMSE was used for comparison (Table S5). To determine whether modules captured outcome-specific variance beyond benchmark scores, we extracted standardized coefficients for module and benchmark terms from the full models for the clearest relationships (A3–tau, A1–amyloid, H6–NeuN; Figure 4C).

#### 4.3.5 Internal and external reproducibility

We examined internal reproducibility by comparing the discovery set with the locked-holdout set. Direction-aligned partial Spearman correlations were calculated for ten Allen readouts and two SEA-AD A3 readouts. For repeated splits, we counted the proportion of results that retained the expected direction. For external RNA analysis, GSE146639 donor scores were modelled with CTR=0, CTR+=1, MCI=1.5, and AD=2 [38], adjusted for age and sex. Equal coding, removal or merging of MCI, and leave-one-donor-out analyses are reported in Table S6. Benjamini–Hochberg correction was applied across the 14 module tests from the final scoring configuration. To test inter-arm independence in the external dataset, we calculated donor-level Harmful and Adaptive total scores and correlated them using Spearman correlation. For external protein replication, we used E-PROT-51 [39]. Protein abundance was transformed as log2(1 + abundance) and standardized. H7 and A2 protein scores were compared among AD, asymptomatic AD, and control groups. Age, an indicator for ages recorded as greater than 90 years, and sex were included as covariates in models with HC3 robust standard errors (Table S6) [40]. Benjamini–Hochberg correction was applied to the four reported protein comparisons.

#### 4.3.6 Single-cell-state basis

For direct reconstruction, each of the six Micro-PVM states was reconstructed using all nuclei from the donor region. Reconstructed scores for each gene group and for the overall arm were calculated using the fraction of each Micro-PVM state and the state-level expression within that state, and were compared with actual scores using Spearman correlation. Because this direct reconstruction reuses the same nuclei, it was treated as a decomposition check. To test whether agreement depended on reusing the same nuclei, we performed a complementary-half reconstruction analysis. Within each donor region, nuclei were randomly split into two halves 100 times. The observed score was calculated in one half, whereas state fractions, state-specific scores, and the reconstructed score were calculated in the other half. The two halves were then exchanged. Cross-half Spearman correlations were calculated across donor regions. The reliability ceiling was defined as the square root of the product of the observed-score split-half correlation and the reconstructed-score split-half correlation. This complementary-half analysis tests reproducibility across non-overlapping nuclei within the same RNA dataset; it is not an external validation of function (Table S7). We also correlated every gene-group score with the intensity inside each state and with the fraction of that state. A donor region was included only when at least 10 nuclei were found in the relevant state. These same-RNA and compositional analyses were used descriptively and were not used to estimate causal state contributions.

#### 4.3.7 Technical robustness

To assess whether donor-level scores depended strongly on cell number, we randomly sampled 200 Micro-PVM nuclei within each eligible donor-region sample and repeated this 100 times. Sampled counts from the same donor were added together before calculating scores. Full-data donor scores were compared with mean donor scores obtained from downsampling. Downsampling was evaluated at the donor level (n = 84) after sampling within donor–region samples. Spearman correlation was used to determine whether rankings were consistent. Bland–Altman bias and limits of agreement were used to determine the difference between them [41]. Single-run ICC(1,1) was used to examine agreement from one downsampling run [42]. For the exact leave-one-gene-out analysis, we removed each of the 94 gene–group entries separately. After one entry was removed, its affected gene-group score and the corresponding overall arm score were recalculated. This process was repeated until every entry had been removed once.

We also calculated scores using rank mean, fixed-reference z-score, robust z-score, GSVA [43], singscore [44], and UCell [45]. Every scoring result was compared with the P10–P90 reference score using Spearman correlation (Table S8). Modules with fewer than four genes (H4, H5, and A6) showed higher deletion sensitivity in leave-one-gene-out analyses. Therefore, interpretation of these specific sub-scores should be module-level only, and their constituent genes should not be used as independent biomarkers without further validation.

## 5. Conclusions

In summary, we developed a function-oriented transcriptional scoring framework comprising seven harmful- and seven adaptive-associated gene modules, designed to resolve terminal microglial functions beyond activation states. This framework addresses the inability of existing state-based scores to determine what microglia do after activation and to compare functional programmes across datasets. Using SEA-AD microglia/PVM nuclei, we computed donor–region pseudobulk scores on a fixed reference scale and validated them against matched protein, activity, and tissue endpoints. Several modules showed expected biological relationships: H3 with cytokine/chemokine proteins, H4 and A5 with oxidative-stress readouts in opposite directions, and A3 with dense-core plaque fraction. Benchmark comparisons indicated partial outcome-specific non-redundancy, and external RNA and protein datasets supported directional reproducibility, although harmful and adaptive arms were highly collinear in the external RNA dataset. The scores were technically stable and reproducible across non-overlapping nuclei at the single-cell level. This framework can be used to compare microglial functional profiles across donors, brain regions, cohorts, and experimental systems, and may help evaluate whether treatments modulate harmful versus adaptive programmes. However, the scores remain observational, are not validated functional measures, and require further mechanistic and prospective validation.

## Supplementary Materials

The following supporting information can be downloaded at https://www.mdpi.com/article/doi/s1, Figure S1-S3: Adjusted residual scatter plots for module–readout relationships; Figure S4: Direct reconstruction of observed scores from microglial states; Table S1: Score configuration: module definitions, gene membership and scoring parameters; Table S2: Cross-cell-type expression attribution of score genes and modules; Table S3: Matched biological readouts and adjusted module-readout associations; Table S4: diagonal specificity; Table S5: Comparison with microglial benchmark scores; Table S6: Internal and external reproducibility analyses; Table S7: Single-cell-state reconstruction and complementary-half reproducibility; Table S8: Technical robustness of harmful and adaptive scores.

## Author Contributions

Conceptualization, methodology, validation, formal analysis, investigation, visualization, writing—original draft preparation, Alex Tu; supervision, project administration, writing—review and editing, S.L. All authors have read and agreed to the published version of the manuscript.

## Funding

This research received no external funding.

## Data Availability Statement

The data analyzed in this study are publicly available. SEA-AD data were obtained from the AD Knowledge Portal under accession number syn26223298. The Aging, Dementia and TBI Study data were obtained from the NIAGADS Data Sharing Service under accession number NG00059. GSE146639 data are available from the Gene Expression Omnibus. The external proteomic data are available from Expression Atlas under accession number E-PROT-51. Statistical results are provided in Tables S1–S8. Supplementary Code lists the inputs required for the included analysis workflows. No new data were generated in this study.

## Acknowledgments

We acknowledge the public data resources used in this study. Part of the data was obtained from the AD Knowledge Portal and the Seattle Alzheimer’s Disease Brain Cell Atlas (SEA-AD). SEA-AD is supported by the National Institute on Aging grant U19AG060909. The SEA-AD data were generated from postmortem brain tissue donated through the University of Washington BioRepository and Integrated Neuropathology (BRaIN) laboratory and Precision Neuropathology Core, with support from the UW Alzheimer’s Disease Research Center, the Adult Changes in Thought Study, prior NIA funding, and the Nancy and Buster Alvord Endowment. We thank the study participants and their families for making these data available for research, and the investigators and staff who maintained and shared these resources.

## Conflicts of Interest

The authors declare no conflicts of interest.

## Notes

### Competing Interest Statement

The authors have declared no competing interest.

